# New high-throughput screening assays for nitrification inhibitors identify an oxazolidine-thione as biological nitrification inhibitor

**DOI:** 10.1101/2024.03.27.586932

**Authors:** Fabian Beeckman, Andrzej Drozdzecki, Alexa De Knijf, Dominique Audenaert, Tom Beeckman, Hans Motte

**Author notes:** **Author contributions** F.B. and A.D.K. performed cultivation and upscaling of the microbial model systems. Griess and Berthelot assays were optimized for use in multi-well plates by F.B. and performed by F.B., A.D.K. and A.D. Method for preservation of microbial cultures was adapted by F.B. and performed by F.B. and A.D.K. Experiments for development of miniaturized nitrification inhibition assays were designed in close collaboration between F.B., H.M., A.D., T.B. and D.A. and performed by F.B., A.D. and A.D.K. Assay development data was analyzed by F.B., A.D. and H.M. High-throughput screening procedures were performed by A.D. with the help of F.B. and A.D.K. The *N. europaea* ammonia vs hydroxylamine oxidation assay was developed by F.B. and executed by F.B. and A.D. The research was supervised by H.M., D.A. and T.B. This manuscript was written by F.B. with the help from H.M., D.A. and T.B.

## Abstract

Nitrification is a microbial two-step process whereby ammonia (NH_3_) is first oxidized into nitrite (NO_2_^-^), by ammonia-oxidizing microorganisms, and then into nitrate (NO_3_^-^) by nitrite-oxidizing bacteria (NOB) in a mutualistic symbiosis. Due to high nitrogen (N) input in agriculture, this process needs to be slowed down as NO_2_^-^ and NO_3_^-^ easily leach, becoming unavailable for plants and leading to eutrophication. Moreover, they may be further processed into the very strong greenhouse gas nitrous oxide (N_2_O). Nitrification thus results in inefficient use of fertilizer and leads to detrimental environmental impact. Inhibitors can be used to mitigate these negative effects and are shown to positively affect plant growth and crop yield and strongly reduce nitrogen (N) pollution. However, currently, only a limited portfolio of nitrification inhibitors is commonly available. These show a variable efficiency and come with logistic complexities. As such, there is a high demand for a broader portfolio of novel molecules. To contribute to the discovery phase of new inhibitors and enable the evaluation of a large number of molecules, we developed miniaturized high-throughput screening assays on the soil-borne ammonia-oxidizing bacteria (AOB) *Nitrosomonas europaea* and *Nitrosospira multiformis*. We give a detailed overview of the procedure and illustrate its use by testing structural variants of oxazolidine-thiones that were previously discovered as nitrification inhibitors. This led to the discovery of the natural compound goitrin as a new biological nitrification inhibitor (BNI). Overall, the presented assays are promising discovery tools, as demonstrated by the newly discovered BNI, and may contribute to the advancement of a more sustainable agriculture.

## 1. Introduction

In agriculture, crops are heavily fertilized with nitrogen (N) to increase yields and ensure sufficient food production. However, fertilizer use efficiency (FUE) is largely reduced by nitrification in soil (Prosser et al., 2020; Lam et al., 2022). The first and rate-limiting step in this microbial process is ammonia (NH_3_) oxidation, which is executed by the ammonia monooxygenase (AMO) enzyme in ammonia-oxidizing microorganisms. Subsequently, nitrite (NO_2_^-^) is formed by the actions of the hydroxylamine dehydrogenase (HAO) enzyme and a yet unidentified component (Caranto and Lancaster, 2017; Beeckman et al., 2018; Beeckman et al., 2023a). Subsequent oxidation of NO_2_^-^ results in the formation of nitrate (NO_3_^-^), serving together with ammonium (NH_4_ ^+^) as a N source for plants (Pélissier et al., 2021). NO_2_ ^-^ and NO_3_ ^-^ are however highly mobile in the soil, and a significant part of it may leach into deeper soil layers. This N loss leads to contamination of groundwater, toxic algal blooms, biodiversity loss (Erisman et al., 2013), and, via downstream processes, to the formation of the ozone-depleting nitric oxide (NO) or highly global warming-inducing nitrous oxide (N_2_O) (Stein, 2019). These nitrogen oxides (NOx) also directly affect human health causing acute or chronic lung diseases (Erisman et al., 2013).

Given that the world population is expected to reach 9 billion people by 2050, intensification of fertilizer use and thus N pollution can be expected, necessitating effective solutions to mitigate N loss. One of the most cost-effective ways to reduce N loss and its pollution is the use of nitrification inhibitors (Winiwarter et al., 2018; Lam et al., 2022). In agriculture, three nitrification inhibitors are commonly used: dicyandiamide (DCD), 3,4-dimethylpyrazole phosphate (DMPP) and nitrapyrin (2-chloro-6-trichloromethylpiridine) (Beeckman et al., 2018). Unfortunately, their efficacy varies with soil characteristics (e.g. pH and moisture content) and climate conditions (Yang et al., 2016; Guardia et al., 2018; Lam et al., 2022; Beeckman et al., 2023a). Moreover, DCD requires high application rates which sometimes leads to phytotoxic effects (Weiske et al., 2001; Wissemeier et al., 2001), while DMPP seems to have a low stability in certain soils (Doran et al., 2018; Sidhu et al., 2021). Finally, nitrapyrin is not registered for use the EU (Papadopoulou et al., 2020). Hence, there is a limited portfolio of nitrification inhibitors whereas it is suggested that repeated use of the same type of inhibitors could lead to resistant soil nitrifying organisms or communities (Subbarao and Searchinger, 2021). These observations underscore the need for novel alternative nitrification inhibitors. In this context, biological materials present an intriguing source, as numerous natural products, often found in root exudates, have demonstrated nitrification inhibitory capacities, termed biological nitrification inhibitors (BNIs) (Subbarao et al., 2009; Coskun et al., 2017; Subbarao and Searchinger, 2021; Lu et al., 2023). BNIs offer potential advantages over synthetic inhibitors, particularly for application (Subbarao and Searchinger, 2021). Many potent BNIs might still be discovered, for instance via screening of fractionated plant extracts or root exudates for nitrification inhibitory activity. Of significant importance, the practical application of new nitrification inhibitors still entails challenging and complex product development, economic aspects, and regulatory processes, often resulting in high fallout rates of candidate inhibitors. Screening efforts can help broaden the pool of nitrification inhibitors to identify those suitable for agricultural applications. However, such screening endeavors, whether using synthetic or natural resources, necessitate the availability of efficient and robust nitrification inhibition assays.

Nitrification inhibition assays typically involve assessing NO_2_^-^ production by ammonia-oxidizing bacteria (AOB) such as *Nitrosomonas europaea* or *Nitrosospira multiformis*, and often use large volumes of bacterial batch cultures (Grunditz and Dalhammar, 2001; Shen et al., 2013; O’Sullivan et al., 2017; Taggert et al., 2021; Yildirim et al., 2022). A modified bioluminescent strain of the AOB *Nitrosomonas europaea* can be used to quantify nitrification inhibition (Iizumi et al., 1998; Subbarao et al., 2006), or colorimetric methods can be employed to measure NO_2_^-^ or ammonium (NH_4_^+^) (Grunditz and Dalhammar, 2001; Duncan et al., 2017; Yildirim et al., 2022). Although these assays typically use a miniaturized set-up in multi-well plates to measure NO_2_^-^ production, the treatments on *N. europaea* or *N. multiformis* occur in test tubes with volumes ranging from 2.5 to 22 mL (Grunditz and Dalhammar, 2001; Duncan et al., 2017). Alternatively, 980 µl aliquots in deep-well plates are used (Yildirim et al., 2022). Hence, all reported assays require relatively high volumes of compounds to be tested, which is undesirable for high-throughput screening (HTS), where thousands of compounds could be tested.

Here, we report on the development and miniaturization into 384-well plate format of two nitrification inhibition assays suitable for HTS using *N. europaea* and *N. multiformis*. We showcase the full procedures of the methods applied during assay development, including guidelines how to maintain and preserve the bacterial cultures. Moreover, we highlight possible pitfalls via multiple examples encountered, which should enable other researchers to establish similar assays. As an example, the *N. europaea* assay was adapted to distinguish between inhibitors of the AMO and HAO enzyme. Finally, to illustrate the applicability of the screening assays, structural variants of oxazolidine-thiones were tested in dose-response, revealing among others the natural compound 5-ethenyl-1,3-oxazolidine-2-thione or goitrin as a new biological nitrification inhibitor. Overall, the implementation of these HTS-compatible assays can contribute to a more efficient discovery of both synthetic and biological nitrification inhibitors.

## 2. Materials & methods

### 2.1 Microbial cultures and cultivation

*Nitrosomonas europaea* DSM28437^T^ and *Nitrosospira multiformis* (NCIMB 11849) were grown in ATCC 2265 and 181 media (NCIMB Ltd), respectively. ATCC 2265 medium was prepared by aseptically combining 900 mL of stock solution 1 (27.75 mM (NH_4_)_2_SO_4_, 3.35 mM KH_2_PO_4_, 0.83 µM MgSO_4_, 0.22 µM CaCl_2_, 11 µM FeSO_4_, 18.33 µM EDTA and 0.56 µM CuSO_4_) with 100 mL of stock solution 2 (400 mM KH_2_PO_4_ and 40 mM NaH_2_PO_4_, pH 8.0 (NaOH)) and 8 mL of stock solution 3 (5 % anhydrous Na_2_CO_3_) solution. All three stock solutions were autoclaved in advance. *N. europaea* cultures were grown at 28 °C and shaken (± 150 rpm) in the dark. Late-log phase cultures ([NO_2_^-^] = 10 - 20 mM, see 2.2. for NO_2_^-^ measurement procedure) were subcultivated by centrifugation (4000 rpm, 15 min, 5 °C) and complete medium refreshment by discarding the supernatant and resuspension of the bacterial cell pellets in fresh growth medium. 181-medium contained per L 1.78 mM (NH_4_)_2_SO_4_, 1.47 mM KH_2_PO_4_, 272 µM CaCl_2_ x 2 H_2_O, 162 µM MgSO_4_ x 7 H_2_O, 1 mL stock solution 1 (1.8 mM FeSO_4_ x 7 H_2_O and 1.49 mM NaEDTA) and 1 mL stock solution 2 (0,5 % phenol red, pH indicator). pH 7.5-8 was maintained by regular additions of sterile 5 % Na_2_CO_3_. *N. multiformis* cultures were incubated in the dark at 30 °C while shaking (± 150 rpm). Late-log phase cultures ([NO_2_^-^] = ± 3 mM) were subcultivated by centrifugation (4000 rpm, 15 min, 5 °C) and complete removal of the old medium and resuspension of the bacterial cell pellets in pH-adjusted 181-medium.

### 2.2 Nitrite measurements

Nitrite was measured via a colorimetric Griess assay (Griess, 1879), by adding 15 µl Griess reagent (Sigma-Aldrich, Cat. No. G4410) to 15 µl culture sample in a flat-bottom 384-well plate (Cat. No. X7001, Low Profile Microplate, Molecular Devices). As the theoretical working range is between 0.43 and 64 μM, it is important to dilute the sample until it is in this working range and calculate the original concentration. For accurate quantitation of NO_2_^−^ levels in experimental samples, a Standard Reference Curve (SRC) was made with NaNO_2_ in the matrix or buffer of the experimental samples using an 8-point 1:2 dilution series starting from 100 μM (incl. a blank sample). After 15 min incubation in the dark at room temperature, absorbance (ABS) was measured at 540 nm using a spectrophotometer (EnVision, Perkin Elmer®).

### 2.3 Culture preservation

Late-log phase cultures (100 mL) were centrifuged in two 50 mL Falcon tubes at 4,000 rpm for 15 min at 5 °C. Supernatant was carefully removed and cell pellets in both Falcon tubes were resuspended in 450 μL fresh growth medium with 50 μL 99.95 % DMSO (= 10 % DMSO). All this was transferred to a 2 mL cryovial and incubated at room temperature during a 30 min equilibration period. Cryovials were then transferred to a Mr. Frosty freezing container and stored at −70 °C. After 24 h, the cryovials could be stored directly at −70 °C. For reconstitution, cryovials were thawed at room temperature and centrifuged at 10.000 g for 15 min at 5 °C. Supernatant was removed and cell pellets were resuspended in fresh growth medium prior to incubation.

### 2.4 Ammonia oxidation vs hydroxylamine oxidation assay

The high-throughput nitrification inhibition assay using *N. europaea* (see 3.2.1) was adapted to be able to discriminate between inhibitors targeting the AMO or HAO enzyme. Different to the standard assay, late-log phase 50 mL *N. europaea* cultures were centrifuged (5,000 rpm, 15 min, 5 °C) and washed 3 times in Falcons tubes with 50 mL in ATCC 2265 medium that was not supplemented with (NH_4_)_2_SO_4_ to remove all remaining N. At the final wash step, 10 mL ATTC 2265 medium without (NH_4_)_2_SO_4_ was added to the Falcons to concentrate the cultures 5 times. Pooled and concentrated cultures were distributed in a 384-well plate with 50 µL per well. As a N-source, 0.5 µL of either 100 mM NH_2_OH or NH_4_^+^ was added to the assay plates as N source (to reach a final N concentration of 1mM). 0.5 µL thiourea (10 mM) and phenylhydrazine hydrochloride (hydrazine, 100 mM) were added to the positive control wells for inhibition of the AMO and HAO enzyme at a final concentration of 100 µM and 1 mM, respectively. As a negative control, 0.5 µL 99.95 % DMSO was added to the negative control wells. Finally, 0.5 µL of the compounds were added to reach the final desired concentration. In this modified assay, the incubation time was reduced to 30 min, directly followed by NO_2_^-^ measurement via the Griess reagent using a 1:2 dilution.

### 2.5 Quantification of assay robustness

A widely used parameter to assess the quality of a HTS assay is Z-prime or Z-factor (Z’). This is a measure to assess whether there is sufficient difference between the positive and negative controls. The Z’ usually varies between 0 and 1, but can be negative as well (see Equation 1). A Z’ of 1 is considered a perfect assay, but in practice should ideally be ≥ 0.5, which indicates a large dynamic range with small data variation. In other words, with a Z’ ≥ 0.5, the window between the positive and negative controls values is sufficiently large for screening, while an assay with a Z’ < 0 shows overlap between positive and negative control values and cannot be used for screening.

### 2.6 Chemicals

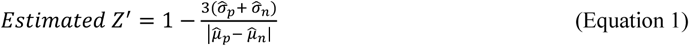

Nitrification inhibitors or candidate inhibitors were acquired from EuroChem (DMP), Sigma-Aldrich (Thiourea, phenylhydrazine, DCD and nitrapyrin), Enamine (3-({[(1,3-benzothiazol-2-yl)methyl](methyl)amino}methyl)-1,3-oxazolidine-2-thione and 2-(methylsulfanyl)-4,5-dihydro-1,3-oxazole), Chemscene ((4S)-4-(propan-2-yl)-1,3-oxazolidine-2-thione, (4S)-4-phenyl-1,3-oxazolidine-2-thione and 5-ethenyl-1,3-oxazolidine-2-thione) or Specs (4,4-dimethyl-1,3-oxazolidine-2-thione).

## 3. Results

### 3.1 Assay plate incubation time

The format in which an assay is performed has a large impact on the behavior of the assay. In a miniaturized format, evaporation can have a large effect on the performance of the assay and this effect is expected to increase with larger incubation times. To test this, we incubated a 96-well plate filled with only 4 mM NaNO_2_^-^ for 72 h and measured the NO_2_^-^ concentration in each well at 0 h, 24 h, 48 h and 72 h. This confirmed increased intra-plate variability over time, especially at 72 h (Fig. 1), and shows that incubation times are preferably kept short, e.g. < 48 h. While technically feasible, incubation periods shorter than 24 hours are undesirable when screening a large number of plates. It is preferable to conduct assay plate preparation, incubation, and read-out simultaneously to prevent overlap of assay steps, which could complicate screening performance.

**Fig. 1:**
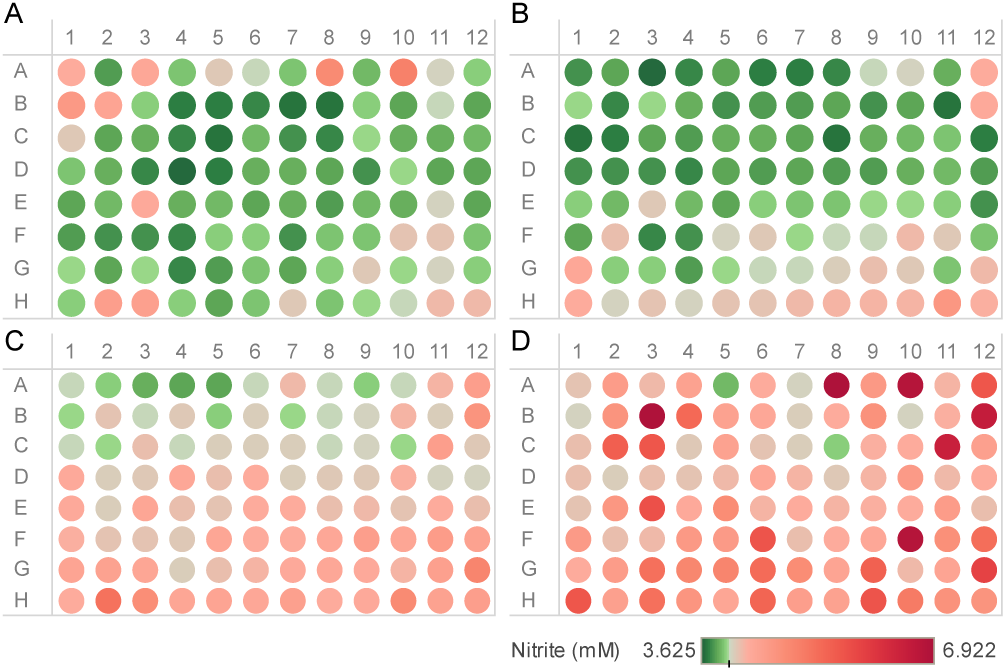
Heatmaps showing the measured NO_2_ ^-^ concentration (mM) in wells of a 96-well plate filled with a fixed 4 mM NO_2_ ^-^ concentration after (A) 0 h, (B) 24 h, (C) 48 h and (D) 72 h incubation. Intra-plate variability and deviation from the initial 4 mM NO_2_ ^-^ concentration clearly increases over time due to evaporation.

### 3.2 High-throughput nitrification inhibition assays

#### 3.2.1 Step-by-step procedure

To perform the high-throughput nitrification inhibition assays, we developed the procedure as shown in Fig. 2. Late-log phase batch cultures of *N. europaea* or *N. multiformis* were distributed in 50 mL Falcon tubes and centrifuged for 15 min at 4,000 rpm at 5 °C. Importantly, late-log phase *N. multiformis* cultures had to reach the late-log phase ([NO_2_^-^] ≥ 3 mM) within at least 3 to 4 days after subcultivation and had to show ≥ 500 µM increase in [NO_2_^-^] over the last 24 h. After centrifugation, the supernatant in all Falcon tubes was carefully poured out in a recipient without disturbing the cell pellet. Remaining supernatant was removed using a micropipette, if needed. Next, the cell pellets were dissolved in 10 mL of the complete growth medium. All dissolved cell pellets were pooled in a sterile Schott bottle before addition to a 384-well plate. The concentrated culture was then used to fill each well of a 384-well plate with 50 µL. As a negative control, 0.5 µL of 99.95% DMSO (or other used solvent for compounds and controls) was added to outer two columns to reach a final concentration of 1%. Similarly, 0.5 µL of 10 mM DMP was added to the other two outer columns to reach a final concentration of 100 µM. DMP was here used to stop nitrification at the time of compound addition and functioned as a reference for the 0h timepoint. Finally, 0.5 µL of 5 mM compounds (or another concentration of stock solution to test other doses) were added to the center wells to reach a final concentration of 50 µM. In this assay, a liquid handling robot Freedom EVO® (Tecan) equipped with a pin tool was used to add the controls and compounds to the assay plates. For this, the pins were washed in sequence with 99.5% DMSO, Milli-Q water and 100% ethanol, and subsequently air-dried in between different additions to the assay plates. To limit evaporation, all plates were individually wrapped with Parafilm and stacked per four on top of a 96-well plate filled with 100 µL Milli-Q per well. These stacks were wrapped in tinfoil and incubated for 24 h at the same conditions as the batch cultures (see 2.1). For the read-out, per assay plate, four U-bottom 96-well plates (= intermediate plates) were filled with 298.5 or 297 µL growth medium per well for a final culture sample dilution of 1:200 or 1:100, respectively. In parallel, per assay plate, one transparent, flat-bottom 384-well plate (= read-out plate) was filled with 15 µL Griess reagent per well. The intermediate plates were used to dilute 1.5 or 3 µL from each well in the assay plates to obtain the necessary dilution for NO_2_^-^ measurement (see 2.2). One intermediate 96-well plate was thus used to dilute the samples of the wells from one quadrant of a 384-well plate. Finally, 15 µL from each well in the intermediate plates was added to 15 µL Griess reagent in the read-out plates. The read-out plates were then incubated for 15 min at room temperature to support the colorimetric reaction. The absorbance in the read-out plates was measured at 540 nm. These ABS540-values were used to finally calculate the relative nitrification using the formula:

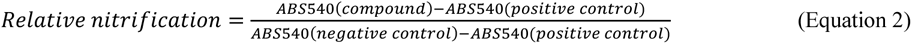

**Fig. 2:**
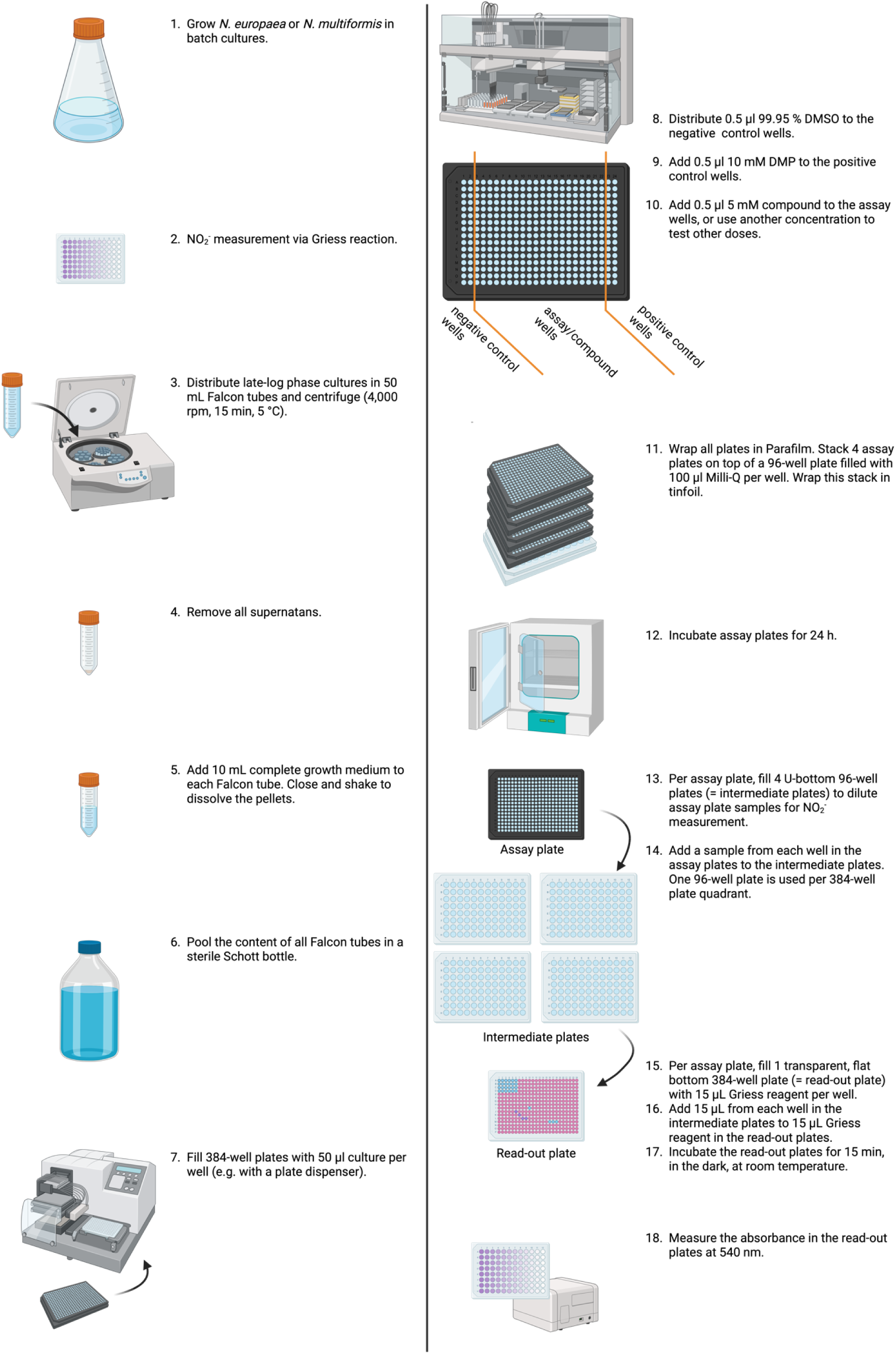
Step-by-step procedure of the high-throughput nitrification inhibition assays. This scheme was created with BioRender.com.

#### 3.2.2 Positive and negative controls for nitrification inhibition

The availability of a positive control is desirable during a HTS campaign as it will allow evaluating the robustness of the assay, measured via the Z-prime (Z’) statistic. In addition, it serves as a reference and allows evaluating the potential difference between a hit and a negative control (no inhibitor). We tested potential positive controls nitrapyrin (100 µM, 20 µM, 4 µM, 0.8 µM), DMP (500 µM, 100 µM, 20 µM, 4 µM) and DCD (2 mM, 1 mM, 500 µM and 100 µM), as well as the negative control DMSO (at 1 %) on *N. multiformis* in 384-well plates and on *N. europaea* in 96-well plates. On both *N. multiformis* and *N. europaea*, nitrapyrin and DMP already showed almost complete nitrification inhibition at 4 µM or 20 µM, respectively, while DCD did not show complete inhibition even at 2 mM (Fig. 3A+B). Surprisingly, in this test the negative controls (1% DMSO and medium) showed high variability in both model systems and, in addition, 2 mM and 100 µM DCD were equally strong. Heatmaps of the read-out plates depicting the absorbance values showed that the wells next to nitrapyrin-containing wells had lower NO_2_^-^ values than expected, which was most clear at the highest concentrations tested (100 and 20 µM) (Fig. 3C-D). This plate effect was thus most likely due to nitrapyrin volatility. The weak inhibition of DCD and the volatility of nitrapyrin makes both inhibitors incompatible for use in a miniaturized assay. We finally validated the inhibitory effect of 100 µM DMP on *N. multiformis* and on *N. europaea* in 384-well plates and found that this consistently showed complete nitrification inhibition (Fig. 3E-F), resulting in a large separation between positive and negative control values, which makes the assay well-suited for HTS. As such, we selected 100 µM DMP as a positive control in both *N. europaea* and *N. multiformis* assays, and 1% DMSO (compound solvent) as a negative control.

**Fig. 3:**
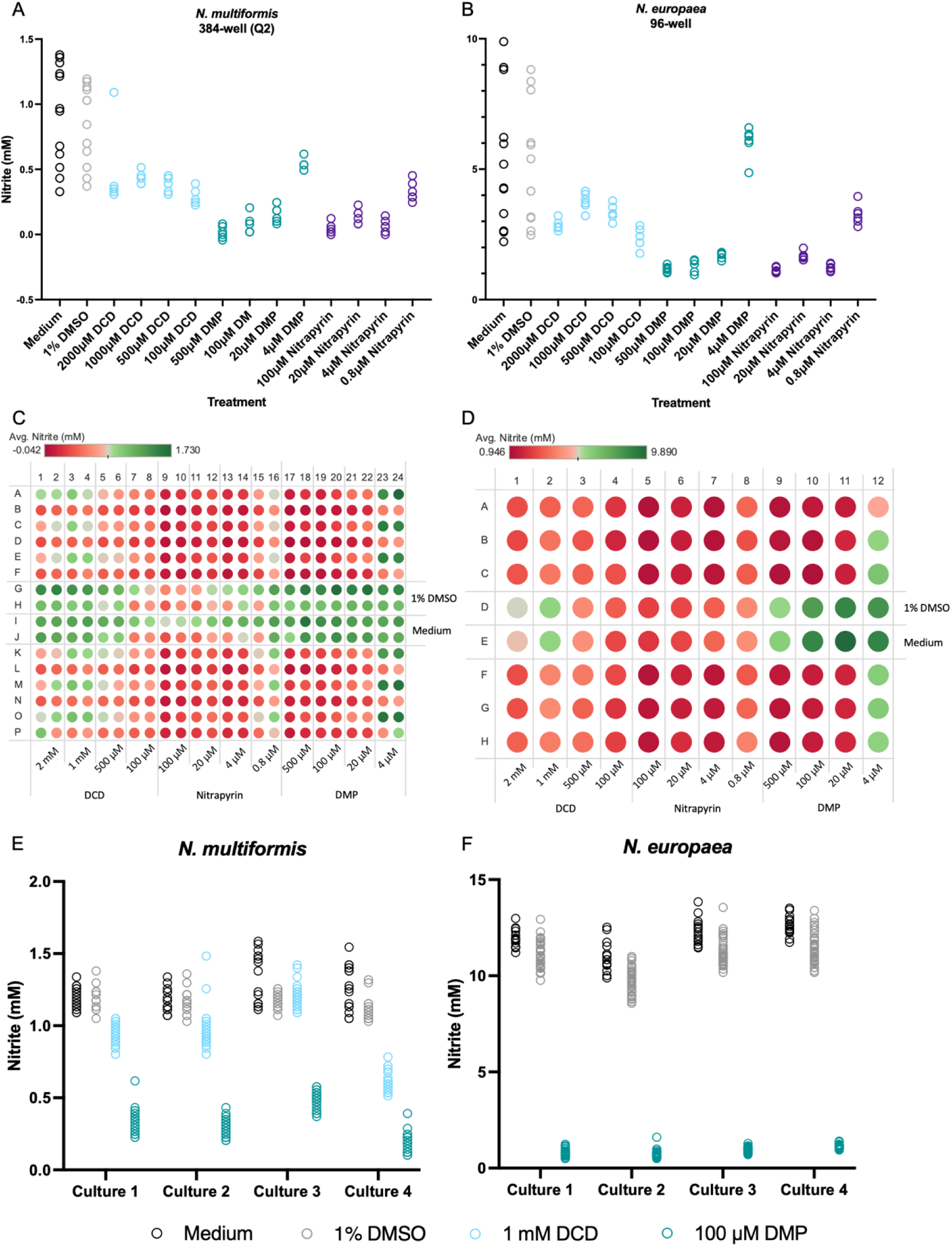
(A-D) Effect of 1% DMSO, DCD (2 mM, 1mM, 500 µM and 100 µM), DMP (500 µM, 100 µM, 20 µM and 4 µM) and nitrapyrin (100 µM, 20 µM, 4 µM and 0.8 µM) on NO_2_ ^-^ production by *N. multiformis* in one quadrant (Q2) of a 384-well plate and NO_2_ ^-^ production by *N. europaea* in a 96-well plate. DMP and nitrapyrin are strong inhibitors at ≤ 20 µM, while DCD is less effective at ≥ 100 µM. Heatmaps illustrate the effect of nitrapyrin volatility on NO_2_ ^-^ read-out in multi-well plates: low DCD concentrations show lower NO_2_ ^-^ values than high DCD concentrations, and wells with 1% DMSO and medium show lower NO_2_ ^-^ values closer to nitrapyrin. (E-F) Effect of 100 µM DMP and/or 1 mM DCD on NO_2_^-^ production by *N. multiformis* and *N. europaea* in 384-well plates. 100 µM DMP consistently showed complete nitrification inhibition on both AOB.

#### 3.2.3 Assay plate preparation time

For a fast and efficient HTS campaign, a large number of assay plates should be run within one batch. The timeframe between addition of fresh growth medium to the bacterial cells and distribution of the compounds to the assay plates (or preparation time) could allow nitrification before compound addition, thereby decreasing differences between positive and negative controls. This can reduce the robustness of the assay, which we measured by the Z’ and ideally should be > 0.5. If compounds were, for example, added to the assay plates shortly after medium refreshment (t = 0 h), very clear differences between positive and negative controls can be observed (Fig. 4), validated by a Z’ of 0.56 ± 0.05 (mean ± SD). This significantly reduces the probability of selecting false positive hits. A similar difference could be observed if there was 3 h in between both handling steps, though the NO_2_^-^ level with DMP treatment increased compared to a 0 h preparation time as nitrification could occur in the 3 h before compound addition (Fig. 4). This resulted in a lower Z’ of 0.48 ± 0.06 (mean + SD). Compound addition 6 h past medium refreshment resulted in multiple samples close to the level of the negative controls just due to NO_2_^-^ produced before compound addition (Fig. 4), with a Z’ of 0.31 ± 0.11 (mean ± SD). These results clearly highlight that assay plate preparation time should be considered carefully, and therefore we opted for < 3 h preparation time. Alternatively, NH_4_^+^ could be added after compound addition, which could be desirable for a small set-up. However, we opted to resuspend the culture in NH_4_^+^-containing medium before distribution of culture, which is for HTS from a practical point desirable as this avoid an extra pipetting step that could increase the risk of technical errors or increase the variability.

**Fig. 4:**
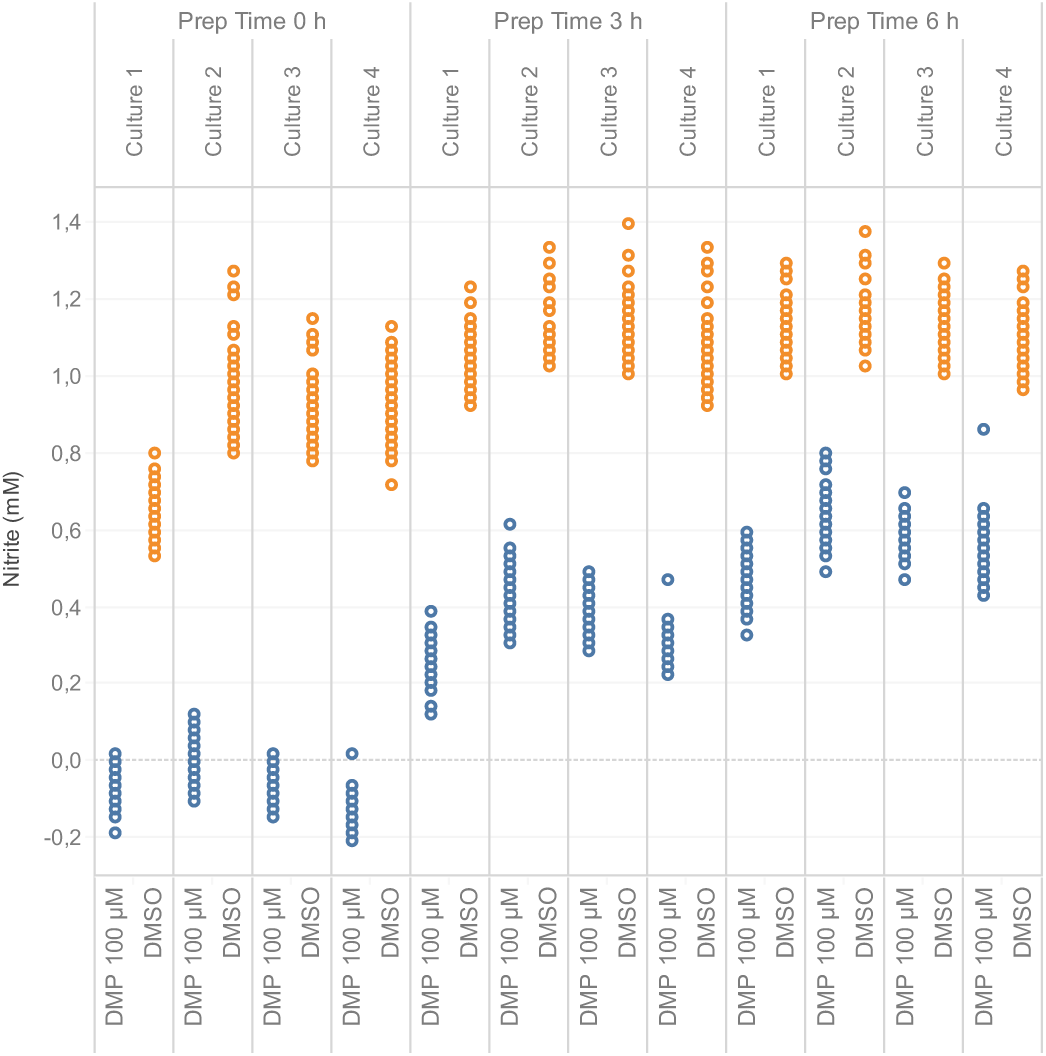
Effect of assay plate preparation time on the *N. multiformis’* assay robustness (Z’ ≥ 0.5), or the difference between positive (DMP) and negative (DMSO) controls. A plate preparation time > 3 h strongly reduces the difference between positive and negative controls, and thus Z’.

#### 3.2.4 Compound screening concentration

Obviously, the concentration of a compound impacts its activity, but when screening high numbers of candidate nitrification inhibitors, it is useful not to use a too high concentration. Besides practical issues such as solubility, high concentrations will make it more difficult to discriminate between strong and weak compounds, while they can also induce artefacts due to precipitation. A too low concentration on the other hand might miss a lot of hits and result in a poor number of hits for further research. To illustrate this, we tested one plate with 320 diverse compounds in the *N. europaea* nitrification assay with a compound concentration of either 10 or 50 µM. To limit the risk of false positives, compounds were selected as hits when the ABS540 value was lower than the ABS540 value of the negative control minus three times the standard deviation of the negative control (Zhang et al., 1999). This revealed 1 hit or 4 hits at 10 and 50 µM screening concentration, respectively, and hence an estimated hit rate of 0.3% or 1.25%, respectively (Fig. 5A). The hit compound in the 10 µM test was also found in the 50 µM test, confirming the robustness of the assay. While the 10 µM test most likely identified the strongest compounds, the hit rate is very low, making 50 µM a more favorable concentration in these assays.

**Fig. 5:**
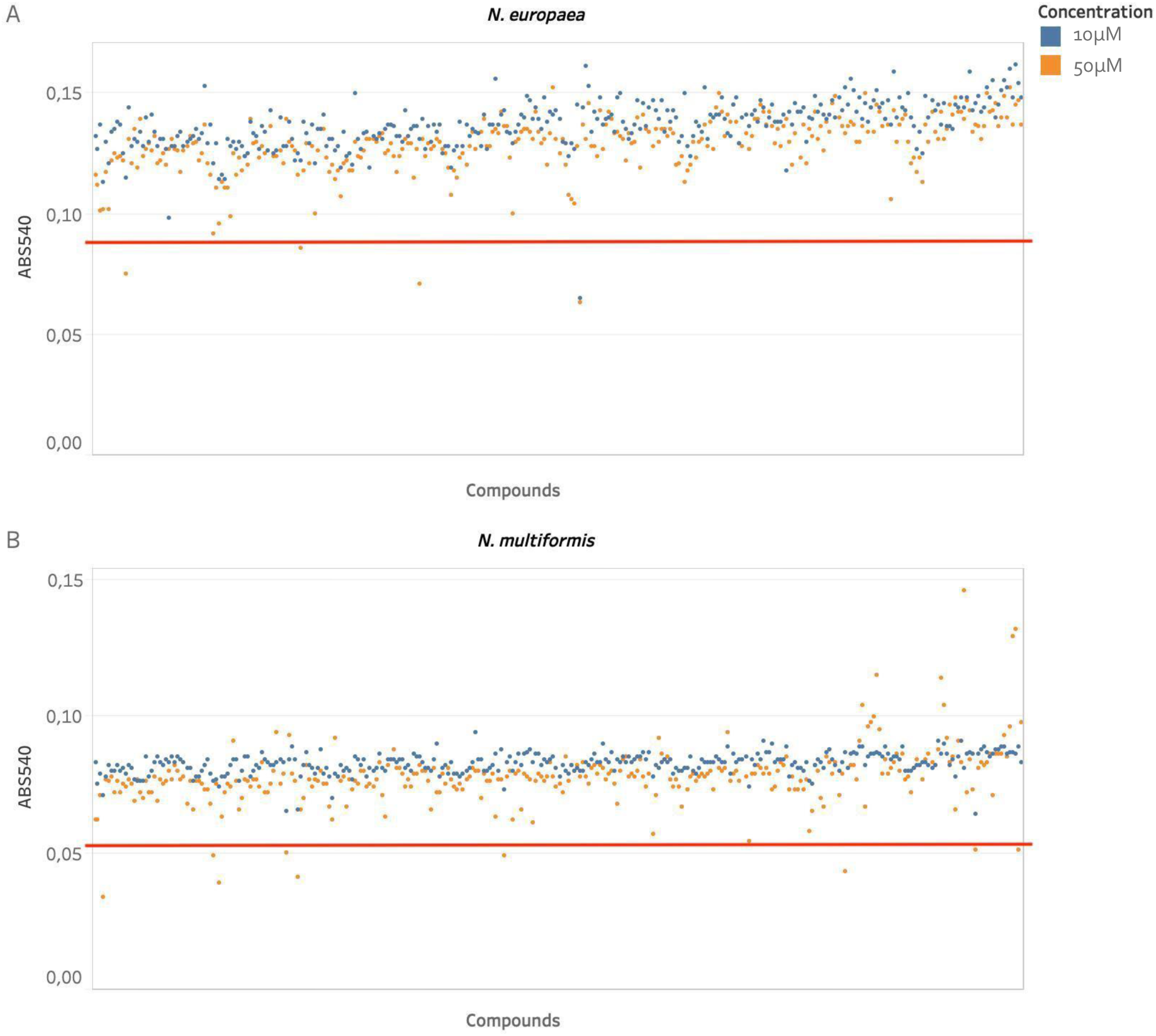
Scatterplots showing theABS540 read-out of a 320 compounds test library screened at 10 µM (in blue) and 50 µM (in orange) on *N. europaea* and *N. multiformis*. The cut-off value (red line) to select the hit compounds (red dots) was set at the ABS540 value minus 3 x SD of the positive control. In *N. europaea* there were 1 and 4 hits at 10 and 50 µM (dots below the red line), respectively, and in *N. multiformis* 0 and 9 hits.

A similar experiment with another test plate was performed with the *N. multiformis* assay. Compounds were again selected as hits as described in the *N. europaea* nitrification assay. This revealed 0 or 9 hits (hit rate of 2.83%) at a concentration of 10 or 50 µM, respectively (Fig. 5B). Therefore, also this assay is preferably performed with compounds at 50 µM.

### 3.3 Ammonia oxidation vs hydroxylamine oxidation assay

To distinguish between inhibitors of the AMO or HAO enzyme, we adapted the HTS assay in *N. europaea*. If the AMO enzyme is specifically inhibited, the inhibitor will still allow the production of NO_2_^-^ if only NH_2_OH is added to the reaction as N source. Indeed, NO_2_^-^ was effectively produced from NH_2_OH if no inhibitor was added. However, DMP appeared to inhibit the NH_2_OH to NO_2_^-^ conversion as well (Fig. 6A). Probably, inhibition of NH_3_ oxidation, crucial to obtain energy, limits growth or the general metabolism of *N. europaea* relatively fast. To rule this out, differences in NO_2_^-^ production were measured at very early timepoints (10, 20, 30 and 60 min after addition of the N source). This showed that both DMP and thiourea directly and continuously blocked the NH_3_ to NH_2_OH step and not the NH_2_OH to NO_2_^-^ step. They could thus be used as positive controls for inhibition of NH_3_ oxidation (Fig. 6B). Notably, the decrease in nitrification rate after 30 min in the negative control (1% DMSO) supports the hypothesis of an indirect inhibition effect through NH_2_OH toxicity at later timepoints (Fig. 6B+C). Finally, phenylhydrazine hydrochloride (hydrazine, 1 mM) immediately blocked NO_2_^-^ production from NH_2_OH and NH_3_, in contrast to DMP or thiourea (at 100 µM) that only blocked oxidation of NH_3_ (Fig. 6C+D). Therefore, it was selected as a positive control for inhibition of NH_2_OH oxidation.

**Fig. 6:**
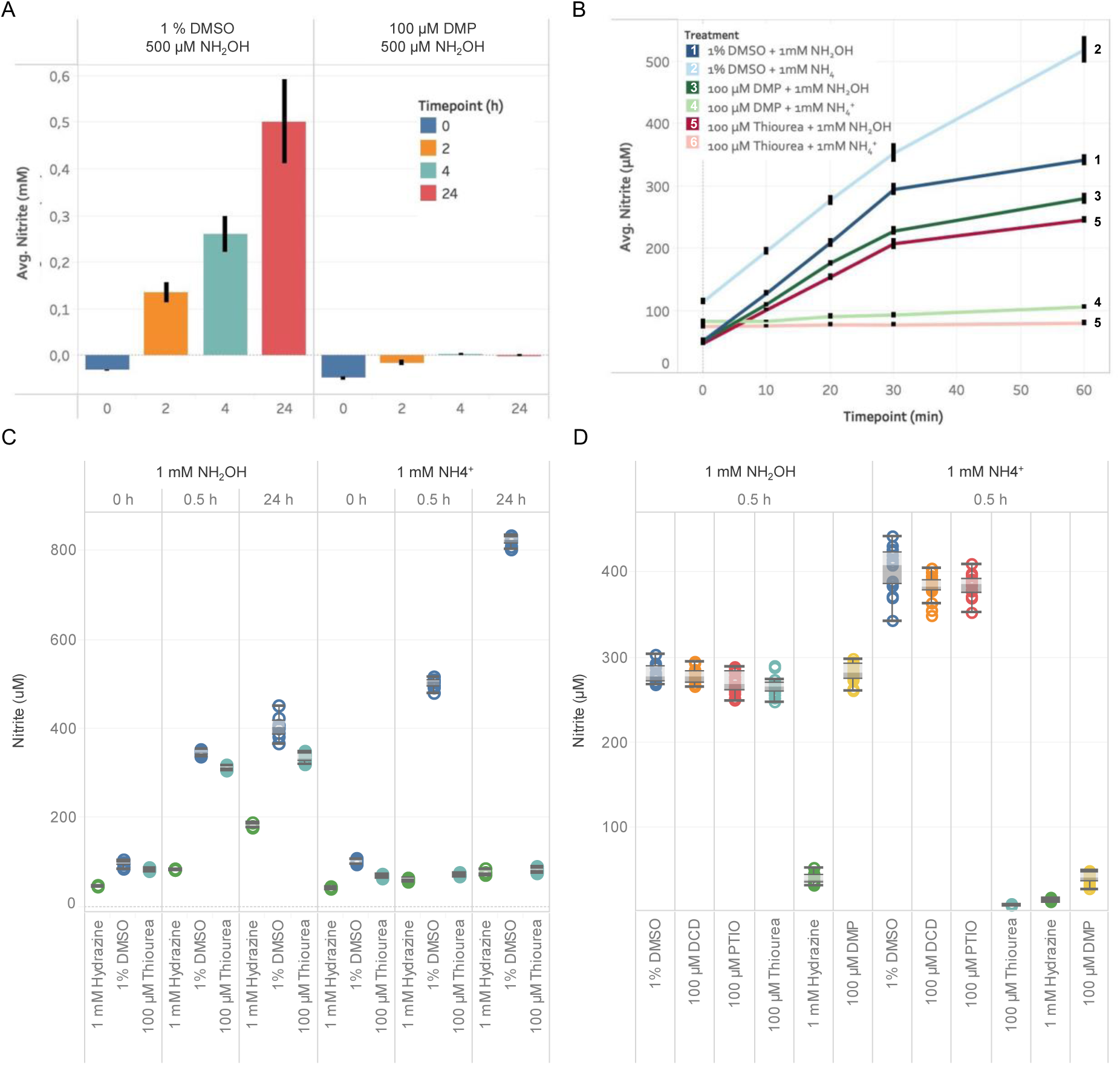
(A) Average NO_2_ ^-^ production from 500 µM NH_2_OH by *N. europaea* at 0 h, 2 h, 4 h and 24 h was blocked by 100 µM DMP, but not by 1% DMSO. (B) Average NO_2_ ^-^ production by *N. europaea* from 1 mM NH_4_ ^+^ or 1 mM NH_2_OH. DMSO, DMP and thiourea treatment (dark blue, green and red, resp.) show increased NO_2_ ^-^ production from NH_2_OH up to 30 min post treatment, in contrast to inhibited NO_2_ ^-^ production from NH_4_ ^+^ by DMP and thiourea. Error bars represent standard deviation (n=8). (C) Boxplots showing the NO_2_ ^-^ production from 1 mM NH_2_OH or 1 mM NH_4_ ^+^ in *N. europaea* cells treated with 1% DMSO, 1 mM hydrazine or 100 µM thiourea. NO_2_ ^-^ production from NH_2_OH stops after 30 min with 1% DMSO, while NH_3_ oxidation continues and is completed at 24 h. Hydrazine has a strong inhibitory effect on both NH_2_OH and NH_3_ oxidation. Thiourea blocks NO_2_ ^-^ production from NH_4_ ^+^, but not from NH_2_OH. (D) Final NH_3_ vs NH_2_OH oxidation assay in *N. europaea*. Thiourea and DMP both blocked NH_3_ oxidation, while NO_2_ ^-^ production from NH_2_OH continued. Hydrazine blocked NO_2_ ^-^ production from both NH_2_OH and NH_4_ ^+^ and is therefore not AMO-specific. DCD and PTIO are no AMO-specific inhibitors under these conditions. Boxplots indicate minimum, 1^st^ quartile, median, 3^rd^ quartile and maximum.

This assay showed that DMP and thiourea immediately blocked NH_3_ oxidation by the AMO enzyme, while DCD, previously reported as an AMO-specific inhibitor, and PTIO did not (Fig. 6D). As such, our original HTS assay proved to be easily adaptable to, for instance, develop an assay for further characterization of nitrification inhibitors.

### 3.4 Testing structural variants of oxazolidine-thiones

We recently discovered a large number of new nitrification inhibitors using the *N. europaea* and *N. multiformis* nitrification inhibition assays, including the strong 1,3-thiazolidine-2-thione (TA-T) (Beeckman et al., 2023b). The substitution of one sulfur atom with an oxygen atom results in 1,3-oxazolidine-2-thione (OAT), a substructure of certain degradation products of the glucosinolate pathway in *Brassicales* species (Agerbirk and Olsen, 2015; Blažević et al., 2020), and which still highly efficient inhibited *N. europaea* as well (Beeckman et al., 2023b). To leverage the *N. europaea* and *N. multiformis* assays, and to further explore inhibitory potential of OAT-containing molecules, we tested a number of variants on *N. europaea* and *N. multiformis*. We acquired several commercially available OAT-containing molecules and tested them in an 8-point dose response experiment in both assays. We used seven different molecules and tested 4 replicates per dose, resulting in 224 samples (excluding controls) per assay. Although the sample size was limited, conducting such experiments in separate flasks would necessitate considerable time and effort, underscoring the utility of the high-throughput assays. As shown in Fig. 7, several OAT-containing molecules, including the natural compound 5-ethenyl-1,3-oxazolidine-2-thione (goitrin), exhibited strong inhibition of both *N. europaea* and *N. multiformis*, comparable to the effect of DMP. A thioether variant of OAT or an isopropyl substitution on position 4 of the OAT ring abolished the inhibitory activity. Other variations retained activity, indicating that other OAT-containing natural molecules could act as BNIs as well.

**Fig. 7:**
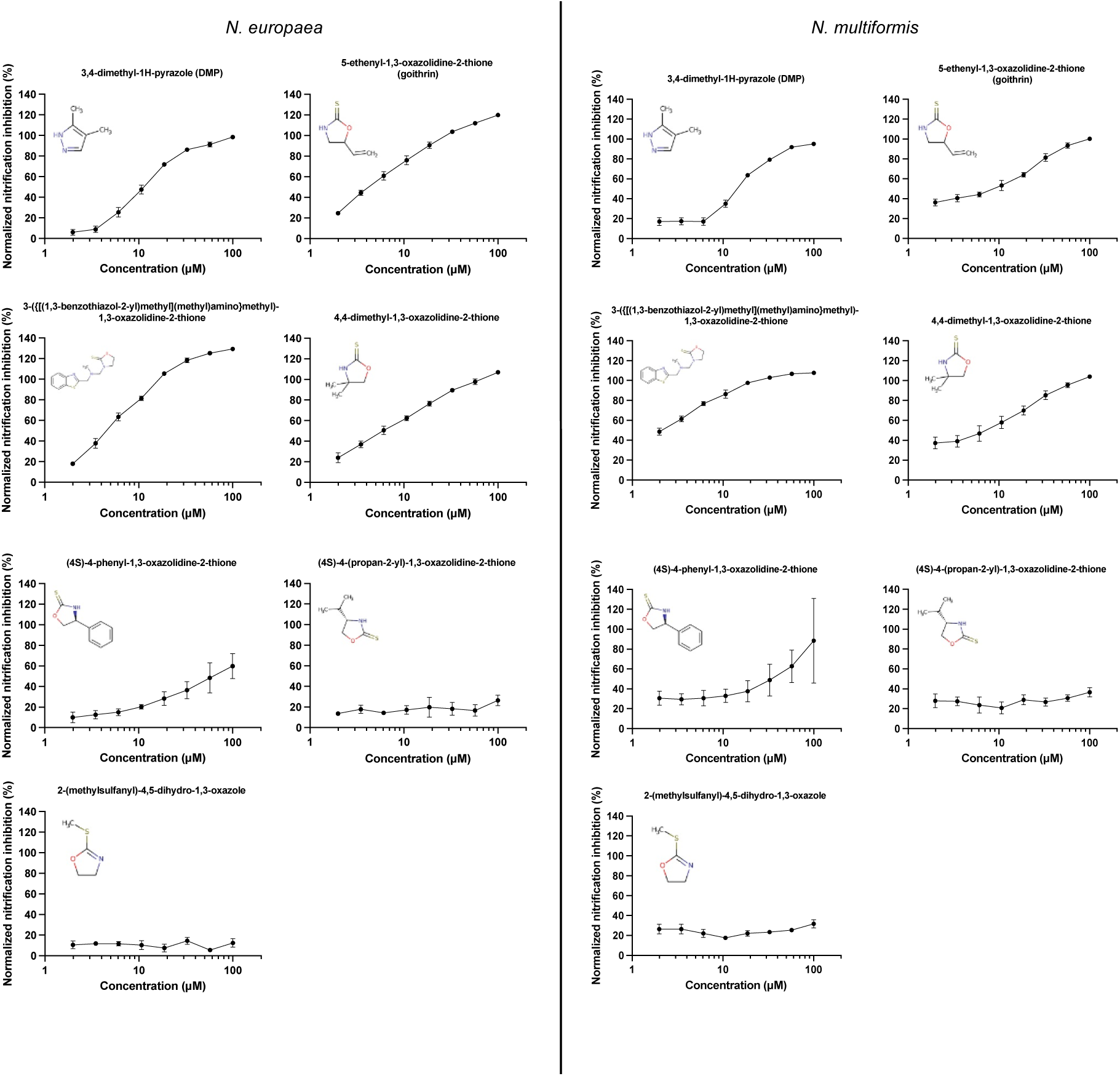
8-point dose response inhibition test on *N. europaea* (left panel) and *N. multiformis* (right panel) with structural variants of oxazolidine-thiones and DMP. Tested concentrations were 100, 57, 33, 19, 11, 6.1, 3.5 and 2.0 µM. The line charts display the average normalized nitrification inhibition percentage (n = 4), and error bars show the standard deviation. Several molecules containing 1,3-oxazolidine-2-thione, incl. the natural compound goitrin show strong inhibition of *N. europaea* and *N. multiformis*.

## 4. Discussion

We here provided detailed procedures for miniaturized colorimetric analyses of nitrification (inhibition) using *N. europaea* and *N. multiformis*. The resulting assays were used for a high-throughput screening campaign that identified new and improved nitrification inhibitors (Beeckman et al., 2023b). Several key factors that can significantly influence the performance of the assays, quantified by the Z-prime (also Z’ or Z-factor), were pinpointed.

Evaporation from multi-well plates during incubation at relatively high temperatures had a profound effect on the Z’ of the assays. Therefore, our tests illustrated that the incubation time must always be limited to 48 h maximum. To assure a sufficiently large window between the positive and negative control in the nitrification inhibition assay and thus a Z’ ≥ 0.5, we aimed for highly active cultures that reached the maximum nitrification inhibition potential before these 48 hours. High activity was obtained by five times concentrating the bacterial cell cultures. Importantly, high nitrification activity limited the assay plate preparation time to max. 3 h. To prevent the cultures from reaching the stationary phase, we chose a 24 h incubation period.

DMSO, the compound solvent, did not negatively influence NO_2_^-^ production by *N. europaea* and *N. multiformis* at 1% using five times concentrated cultures. As such, compounds could still be added to 384-well assay plates, filled with max. 50 µl per well, via a pin-tool that has a fixed working volume of 0.5 µl. Alternatively, it would either require a more costly acoustic dispenser that allows for addition of nanoliter volumes of the compounds, a pre-dilution step of the compound stock solution, or the use of 96-well assay plates.

To find a good positive control for the screening assays, enabling complete inhibition of nitrification at the start of the treatments, multiple nitrification inhibitors were tested at different concentrations. Different (high) concentrations of DCD were first tested as positive controls, as DCD was previously reported to fully inhibit nitrification by *N. multiformis* at 100 µM (EC_50_ = 80.28 ± 6.20 µM) (Shen et al., 2013) and to be 4-fold more effective against *N. europaea*, with an EC_50_ of 119 and 460 µM, respectively (O’Sullivan et al., 2017). However, even at concentrations up to 2 mM, DCD was unable to inhibit nitrification efficiently and consistently in *N. europaea* and *N. multiformis*. Alternatively, 100 µM DMP completely inhibited nitrification in both model systems, and although nitrapyrin required a lower dose for the same effect, it was not selected as a positive control due to volatility. After all, it has been described as a volatile compound (Redemann et al., 1964; Bédard and Knowles, 1989; Ruser and Schulz, 2015), which was clearly demonstrated here as it negatively influenced NO_2_^-^ production in surrounding wells. This displays another point of attention: in contrast to experiments in recipients with bigger volumes where volatility is no issue, volatility of compounds needs to be seriously considered in multi-well assays as it may confound results.

The last step in assay development was the selection of the compound screening concentration. Therefore, a small set of 320 random compounds was tested to assess the hit rate at different concentrations. Ideally, the hit rate should be around 1 to 2%, which could in our case be obtained at 50 µM, and which we previously used to screen 45,000 compounds of a small molecule library on *N. europaea* and *N. multiformis* (Beeckman et al., 2023b). Depending on the resource, it could be recommended to perform a similar test at different doses or dilutions. The ideal dilution of (fractionated) plant extracts, for example, may be affected by the extraction conditions or size of the fractions.

Our newly developed HTS assay on *N. europaea* was used to develop a miniaturized assay that discriminates between compounds working on NH_3_ or NH_2_OH oxidation. This would provide more insights into the possible mode-of-action of (new) nitrification inhibitors. Most current nitrification inhibitors are described as Cu-chelators, potentially binding Cu^2+^ in the active site of the AMO enzyme, thereby preventing the binding of NH_4_^+^ in this site (Beeckman et al., 2018). In our AMO-specificity assay we tested both DMP and thiourea, an inhibitor that is better documented to be AMO-specific, including reports describing similar assays as the one we applied (Subbarao et al., 2006). Hydrazine, previously described as a specific inhibitor of the HAO enzyme (Logan and Hooper, 1995), was selected as a positive control for inhibition of NH_2_OH oxidation. This showed that DMP and thiourea were strong inhibitors of NH_3_ oxidation by the AMO enzyme. In contrast, DCD, that was previously reported as an AMO-specific inhibitor (Zacherl and Amberger, 1990; Lehtovirta-Morley et al., 2013; Shen et al., 2013; Sun et al., 2016), and PTIO did not. The latter was earlier described as a NO-scavenger with a strong inhibitory effect on AOA (Yan et al., 2012; Shen et al., 2013; Martens-Habbena et al., 2015; Sauder et al., 2016). Finally, this derived assay displays the robustness and versatility of the originally developed high-throughput nitrification inhibition assays.

In the end, the development of these assays resulted in detailed procedures of two miniaturized nitrification inhibition assays in two soil-borne AOB. To our knowledge, this has never been described, and ultimately validated (we refer to Beeckman et al., 2023b), in such an extensive manner before. The different discussed pitfalls in the assay development, as well as the detailed high-throughput procedures form a guideline and basis to develop and perform new similar screening assays or efficiently conduct nitrification inhibition experiments. This will be important as there is a clear need for more and diverse nitrification inhibitors to increase nutrient use efficiency in agriculture. To illustrate the possible utilization of the assays, we tested the inhibitory activity of a number of OAT-containing molecules and showed that the naturally occurring goitrin is a novel BNI. Goitrin and other OAT-containing molecules are degradation products of the glucosinolate pathway, which are found in *Brassicales* species (Blažević et al., 2020). These species naturally inhibit nitrification (Brown and Morra, 2009), which is so far attributed to isothiocyanates (Bending and Lincoln, 2000; Brown and Morra, 2009; Balvert et al., 2017). Isothiocyanates are just like OAT-containing molecules derived from the glucosinolate pathway (Blažević et al., 2020). By leveraging the presented high-throughput assays, we here showed that, in addition to the isothiocyanates, also OAT-based degradation products of the glucosinolate pathways may add to the natural nitrification inhibition by *Brassicales* species, and that OATs could be classified as new BNIs.

In conclusion, we showed that our approach opens new avenues for identifying novel nitrification inhibitors. Therefore, we anticipate that our research may play a pivotal role in advancing eco-friendly strategies for managing nitrification and increasing nutrient utilization in agricultural practices.

## Acknowledgements

We thank EuroChem Agro to financially support this research and Peter Bogaert for technical support.

